# Phenotypes in insect biodiversity research

**DOI:** 10.1101/032425

**Authors:** István Mikó, Andrew R. Deans

## Introduction

Phenotypes, here defined as the observable characteristics of organisms, including morphology, behavior, and physiological products (*e.g*., nests and galls)—the results of gene expression, in an environment—are the primary data for research on the evolution and taxonomy of insects (Deans *et al*., 2012a). The shapes, textures, colors, or simply the presence or absence of anatomical features or behaviors help us determine taxa and guide our hypotheses regarding their evolutionary history. Phenotype data are relevant to most other research on insects as well, including biomechanics (Labonte and Federle, 2015), bioinspired engineering (Werfel *et al*., 2014; Lau *et al*., 2014; Winegard *et al*., 2014), genotype-phenotype associations (Houle and Fierst, 2013), genetic modification (Wang and Jacobs-Lorena, 2013), human disease research using insect models (Washington *et al*., 2009), plant-insect interactions (Whitney and Federle, 2013), and many others.

Despite their broad relevance, phenotype data generally lack the standards necessary for computation or even broad understanding by humans. That is, one cannot simply query across phenotype data (What are the most frequently described color phenotypes for Hymenoptera?) with the aid of computers, as one can do with molecular data (What are the most frequently sequenced protein-encoding genes for Hymenoptera?). Researchers increasingly recognize that lifting this barrier will result in more meaningful, integrative research across the life sciences (Deans *et al*., 2015), and computational mechanisms are increasingly implemented in biodiversity research.

## Phenotype data: past and present

Phenotypes have long been the primary data for classifying biodiversity. Even a casual reading of Aristotle’s *History of Animals*, arguably the first hierarchical, scientific classification of life, reveals rich descriptions of arthropod phenotypes. This description of a decapod, for example, reads almost like a modern morphological treatment (Barnes, 1984, Book IV: 526a1–526a12; Aristotle, translated by d’A. W. Thompson):

> In the crayfish the male differs from the female: in the female the first foot is bifurcate, in the male it is undivided; the belly-fins in the female are large and overlapping on the neck, while in the male they are smaller and do not overlap; and, further, on the last feet of the male there are spur-like projections, large and sharp, which in the female are small and smooth. Both male and female have two antennae in front of the eyes, large and rough, and other antennae underneath, small and smooth. The eyes of all these creatures are hard, and can move either to the inner or to the outer side. The eyes of most crabs can do the same, to an even greater degree.

Biodiversity researchers continue to publish prosaic descriptions like this, including hypotheses of function and evolutionary relevance, but most taxonomic descriptions are written in a quasi-standardized, telegraphic style (Mayr and Ashlock, 1991). Consider this descriptive statement of a bee species, also from Aristotle (Barnes, 1984, Book IX: 627b23-971): “another kind is what is called the robber-bee, black and flat-bellied”. In a contemporary taxonomic context, these characters would be written as:

Body black. Metasoma flattened.

Although this simplified prose is arguably more concise and accessible than full sentences, phenotype annotations (data) continue to be generated using personalized lexicons, often impossible to reference or reconstruct, and without a truly standard syntax. Synonymy and homonymy are rampant (Seltmann *et al*., 2013; Yoder *et al*., 2010), and the interpretation of descriptions requires significant human brain power. One could express the above phenotype, that the body is black and the metasoma is flattened, myriad ways:

– Body black, abdomen flattened
– Blackish, with dorsoventrally compressed metasoma
– Fully melanized; posterior tagma flat

At this point in time, only a human would consistently recognize these as roughly equivalent statements. And only a human reader could look past the lack of precision or outright inaccurateness of these descriptions. The entire body, for example, is rarely uniform in color. The parasitoid wasp in Figure 1 was described by Dodd (1920) as being “black” (a few sclerites excepted, including the legs and scape). Yet cuticle of the wing blade and compound eye, each of which is logically part of the body, is transparent. Other parts of the body—ocelli, processes, articulations, setae—likewise are not black, though a reader could assume they are, based on this description. Despite the imprecision of these annotations, most scientists could read them and understand that the body, and most likely the external cuticle alone, is mostly black.

**Figure 1:**
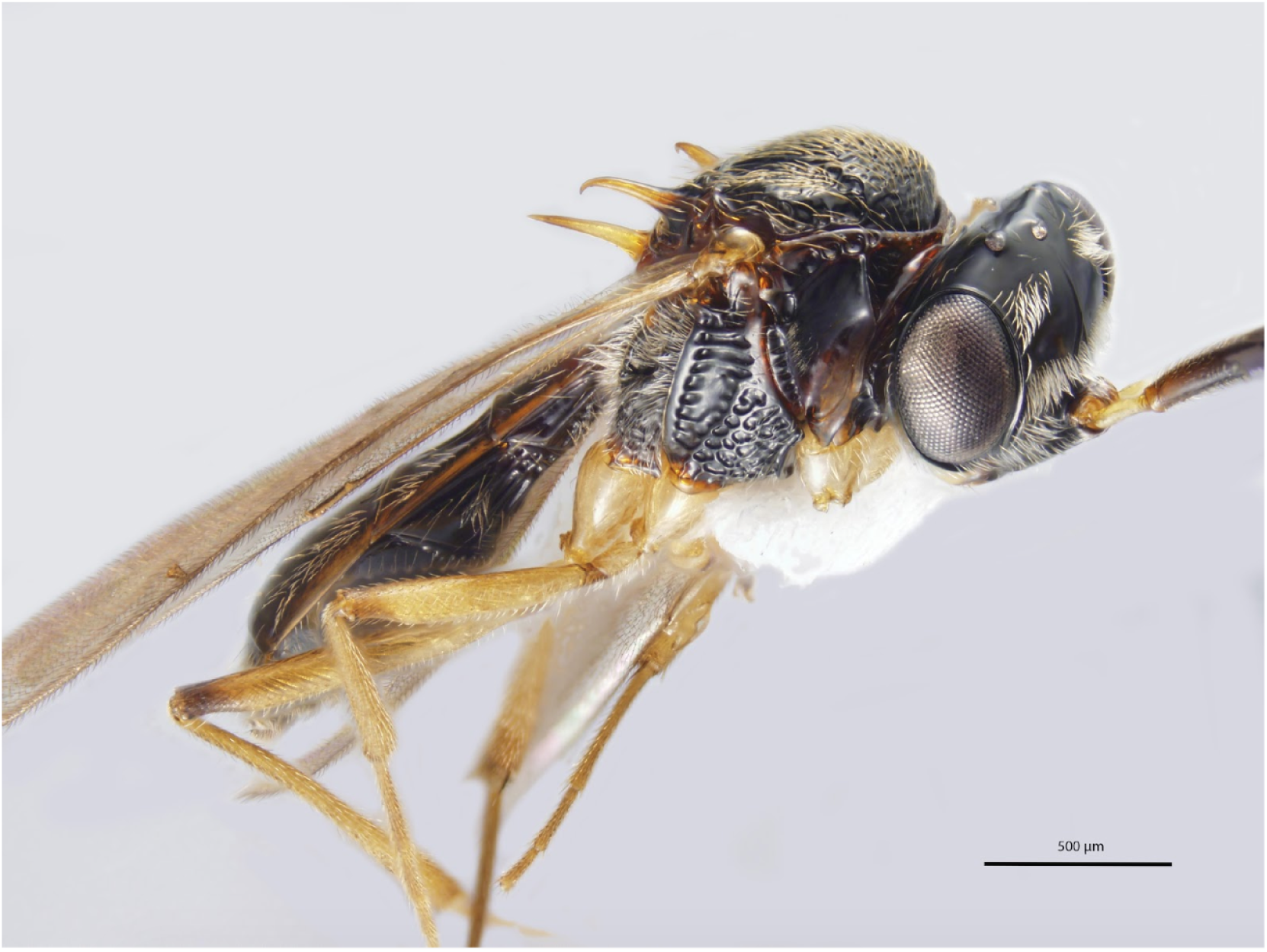
Bright field micrograph of *Gryonoides glabriceps*, originally described by Dodd (1920) as being, in part: “Black; thorax and base of abdomen slightly suffused reddish; leg wholly yellow; antenna1 scape yellow”.

The broader scientific enterprise, however, depends on accuracy, especially when analysis relies on computers. For phenotype data generated by taxonomists to be re-purposed outside of taxonomy, *i.e*., beyond a small community of highly trained domain experts, those data must be generated and availed in a more accessible and meaningful form.

## Phenotype data: present and future

Taxonomists are not the only scientists who generate phenotype description in natural language. Despite some calls for a shift towards representing phenotypes as images, rather than text (MacLeod *et al*., 2010; Riedel *et al*., 2013), the practice is pervasive throughout the life sciences. Communities that work on model species recognized early on that computable phenotypes would increase the potential for novel discoveries (Washington *et al*., 2009, *e.g*.), and they have developed several approaches to make phenotype data more accessible. Most of these systems are built around **ontologies**, which increasingly facilitate biodiversity research as well.

### Biological ontologies

An ontology, at least in this context, is a formal representation of concepts, usually referred to as **classes**, in a particular **domain**, and the **relationships** between those concepts. One can think of an ontology as “a set of well-defined terms with well-defined relationships” (Ashburner *et al*., 2000). Consider this entry from a well-known informal ontology, the *Torre-Bueno Glossary of Entomology* (Nichols, 1989, pg. 654; edited here for simplicity):

> **scape, scapus**; shaft (R. W. Brown); the first or basal segment of the antenna

The definition (the first or basal segment of the antenna) is a class in the domain of entomology, which most researchers refer to by the **terms** “scape” or “scapus” (except R.W. Brown, who uses “shaft”; see discussion of *sensu*, below). The eight-word definition also contains information regarding the relationships of this class to other entomological classes. One can understand that this class is a type of, or **subclass** of, “segment”, and it has the **property** of being *part of* the “antenna”. After reading the definition of segment (“subdivision of … an appendage … associated with muscle attachments” (Nichols, 1989, pg. 667)) the reader can then deduce that because the scape is a type of segment it must be associated with muscle attachments.

Because of the way it is written the substantial knowledge represented in the *Torre-Bueno Glossary of Entomology* is only available, at least in a meaningful way, to humans. A computer could not pick up a paper copy of the book, flip to the definitions of scape and segment, and logically conclude that scapes must have muscle attachments. Classes and their relationships must be **formalized**—*i.e*., written in a form that computers can process—in order to facilitate machine-based computation across this kind of data.

Only a handful of formal ontologies exist for insect biodiversity research. The Hymenoptera Anatomy Ontology (HAO) (Yoder *et al*., 2010) is the most well-developed multi-species example for arthropod anatomy, and several others exist in other domains relevant to phenotypes, specimens, *etc*. (Table 1). Formal ontologies can be developed using a number of approaches, but the most widely used standard in biology is Web Ontology Language (OWL; World Wide Web Consortium (2015)). An OWL representation of an ontology allows one to compute across phenotype data sets in order to address scientific questions and infer relationships that are not explicitly stated (see Figure 2).

**Table 1:** Some ontologies relevant to insect biodiversity research. Note that several other ontologies are available and/or required for proper formalization of insect phenotype data.

| Ontology | Description | Reference |
| --- | --- | --- |
| Hymenoptera Anatomy Ontology | Used for descriptive taxonomy and comparative morphology of Hymenoptera | Yoder *et al.* (2010) |
| PATO | Ontology of phenotype descriptors, used in multiple research domains | Mungall *et al.* (2010) |
| Biospatial Ontology (BSPO) | Biospatial terms, like dorsal, ventral, *etc.* | Mungall (2015) |
| Environment Ontology (ENVO) | Includes classes relevant to environment and habitat | Buttigieg *et al.* (2013) |
| Biological Collections Ontology | Includes classes relevant to natural history collections | Walls *et al.* (2014) |
| Gene Ontology (GO) | Probably the most widely used ontology the life sciences; covers genes, their products, and processes | Ashburner *et al.* (2000) |

**Figure 2:**
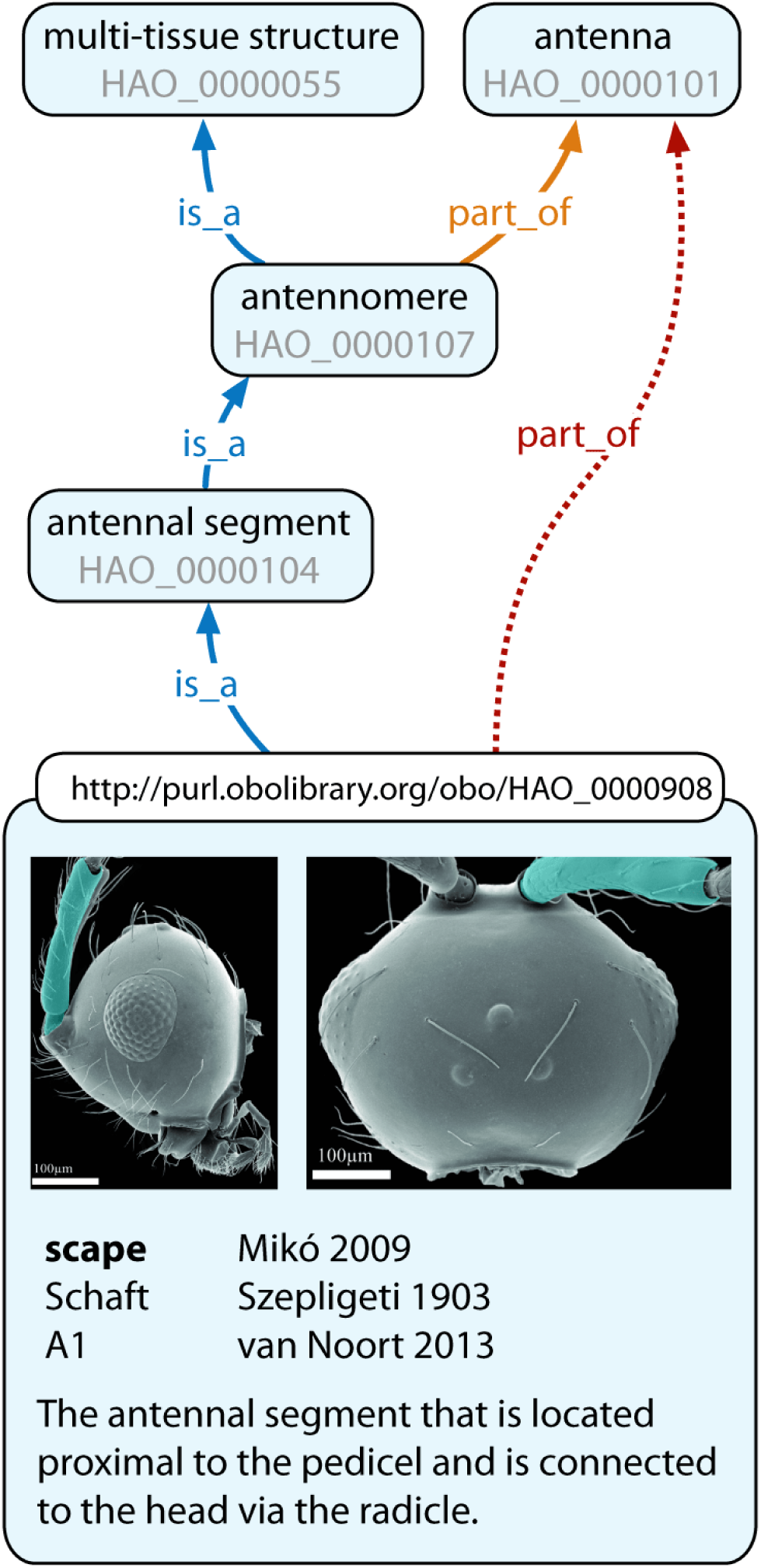
Hymenoptera Anatomy Ontology (HAO). Partial graph of the HAO, illustrating the explicitly stated properties of the scape. The curators of this ontology explicitly relate (solid arrows) scape to antennal segment, using the “is a” relationship. There is no explicitly stated “part of” relationship for scape, but one can *infer*, through properties of anatomical entities higher in the hierarchy, that the scape is part of the antenna (dashed arrow). Scape itself is expanded to reveal its URI (resolvable identifier, at top), instances (annotated images), labels used in the literature (bold name is preferred term), and *genus differentia* definition.

### Ontologies in biodiversity research

Insect biodiversity scientists have yet to integrate large-scale, machine-based computation into their research on phenotypes, but the potential for discovery is substantial. Taxonomic and other morphological descriptions are unparalleled sources of information about evolutionary novelties and adaptations (Deans *et al*., 2012a,b), which, as discussed above, remain hidden from most researchers. The following use cases, some of which are simple to implement, serve as examples where data generation and the process of discovery could benefit from an ontological approach.

**Referencing a glossary** While formal ontologies are primarily designed for machine-based computation, the process of developing these ontologies and the development of user-friendly tools can facilitate human-based computation as well. In a recent analysis of 428 arthropod systematics papers, Deans *et al*. (2012a) found that fewer than 3% of authors included a glossary of morphology terms used in character descriptions. Almost one third of the papers did not cite any references for term meanings. This situation is problematic, especially when terms frequently refer to more than one concept (*e.g*., “paramere” can mean at least five different body parts, and the “forearm” of Lepidoptera is quite different from the forearms of Mammalia). With community-selected “preferred terms” for classes, an ontology can function as a controlled vocabulary, which unifies within and cross-domain knowledge (Noy and McGuiness, 2001; Smith *et al*., 2007) and improves interdomain communication. Even without preferred terms, an ontology can play a central role in word sense disambiguation of anatomical terminology, for example as an online glossary. An online description, marked up with Web links (URLs) that reference explicit concepts can provide more information that can not be stored in the ontology itself, *e.g*., images.

For hymenopterists (and really, the tools would work for most insect systematists), there are tools (Seltmann *et al*., 2012) that match natural language descriptions to concepts in the HAO, so that a table of terms, definitions, and URIs (in this case, links to Web pages with more information) can be generated.

**Generating logically consistent phenotypes** Appending a glossary of logical, structured morphology definitions to one’s manuscript should increase the accessibility of those data, at least from a human perspective. Adding logical, structured *phenotype* descriptions is the next step in making one’s data available for computation. Composing structured phenotype descriptions also, in our experience, improves character development by forcing the researcher to think carefully about the phenotypes being described. Is the character overly complex? Are the states consistent and relevant?

Consider the color of the scape of the wasp in Figure 1. One could create a character called “scape color”, with the states: (0) scape brown and (1) scape black. Brown and black are both children of color in the Phenotypic Quality Ontology (PATO), and therefore these states are perhaps more likely to be homologous. Attempts to add inconsistent character states, for example (2) scape hairy and (3) scape rugose, would stand out as inappropriate. Hairy is a child of pilosity, and rugose is a child of texture; the states are unlikely to be homologous (Although homology can be messy!). The example is a bit silly, perhaps, but problematic character states are pervasive in insect systematics. This character from Whiting *et al*. (1997), edited for clarity, is a bit more subtle: Ovipositor (0) absent, (1) present, (2) vestigial, (3) modified (reduction in the second valvulae, the third valvulae serving as the functional components of the ovipositor), (4) fused. The character combines phenotypes that are related to count (presence, absence), structure (fused, vestigial), and morphology (modified). Are these states homologous, or should the phenotypes be scored as separate characters? Another character from this data set suffers from similar problems: Trochantin (0) absent, (1) present, (2) trochantin-episternal sulcus present. To a naive reader, this character appears to refer to counts of two different anatomical entities (*i.e*., the states are not homologous), the trochantin itself and a sulcus that is associated with the trochantin. We are not suggesting that these characters are not phylogenetically relevant, but the way that the states are textualized make the phenotypes difficult to understand and therefore inaccessible.

In our own research we found many examples where character states were too loosely described to be meaningful, and a semantic approach improved their description. Mullins *et al*. (2012) and Balhoff *et al*. (2013) provide examples of biodiversity research that incorporate explicit, logical representations of phenotypes, composed in OWL using many of the ontologies in Table 1. Mikó *et al*. (2015) provide details about how to generate data in this format, examples of which are in Table 2.

**Table 2:** Typical character statements and their semantic representations.

| character | semantic annotation |
| --- | --- |
| Antennal shelf count: present | has part some antennal shelf |
| Anterolateral mesopectal projection 2-d shape: square shape | has part some (anterolateral mesopectal projection and (has part some (lateral side and (bearer of some square)))) |
| Body length universal: 4.6–8.0 mm | has part some (body and (has part some (median anatomical line and (bearer of some (length and (is quality measured as some ((has measurement unit label value millimeter) and (has measurement value some (float[≥4.6f] and float[≤8.0f])))))))) |
| Male petiole length *vs.* width: $2.5 \times$ as long as wide | has part some (petiole length and (bearer of some (length and (is quality measured as some ((has measurement unit label some (width and (inheres in some abdominal segment 2))) and (has measurement value value 2.5)))))) |
| Mandibular teeth count: 4 | has part some (mandible and (has component exactly 4 tooth)) |
| Ventro-lateral region of mesosoma texture: areolate | has part some (mesosoma and (has part some (ventro-lateral region and (bearer of some areolate)))) |

**Reasoning across phenotype data** When phenotype data are generated logically, as described above, using multiple ontologies and standards of the Semantic Web, one should be able to use the logic inherent in those ontologies for computation. Numerous real and hopeful examples of computation have been published, and they illustrate a range of possible use cases, including image management (Ramírez *et al*., 2007), connecting human disease phenotypes to mutant model organisms (Washington *et al*., 2009), species determination through successive queries (Deans *et al*., 2012b), enhancing taxonomic practice (Balhoff *et al*., 2013), understanding synapomorphies and homoplasy (Ramírez and Michalik, 2014), ontology-based partitioning for phylogenetic analysis (Tarasov and Génier, 2015), linking morphometric data to descriptions (Csösz *et al*., 2015), correlating phenotypes with environment (Thessen *et al*., 2015), a similarity-based approach for recognizing comparative homologs (Vogt, 2015), and correlating phenotypes with genetics and environment, across multiple disciplines (Deans *et al*., 2015).

Prospecting for relevant non-model species is another process that could be facilitated by a semantic phenotype approach. Genomic and developmental approaches continue to grow, with an extraordinary amount of research done on insect models, like *Drosophila* and *Tribolium*. Each of these systems, coincidentally, has its own phenotype and anatomy ontologies (Osumi-Sutherland *et al*., 2013; Dönitz *et al*., 2013), but are they the most well-suited model species for research on certain phenotypes? Could vision impairment in humans be better understood by studying lineages with natural variation in eye development? If one could query across known insect phenotypes—show me all the insects that exhibit eye reduction—other models might reveal themselves and perhaps facilitate compelling comparisons. Research on other developmental processes, *e.g*., branching morphogenesis or epithelial folding, likewise could benefit from increased accessibility to phenotype data from the natural world (see Figure 2 in Deans *et al*., 2015).

Workflows are now available for generating computable phenotype annotations, using either manual or automated approaches. Human-mediated annotations, using applications like Phenex (Balhoff *et al*., 2010) or Protégé, yield relatively precise and accurate data, but the approach is not scaleable to the vast corpus of legacy phenotype treatments. Formulating semantic statements currently requires some knowledge about OWL ontologies and Manchester syntax (Table 2) and likely requires a more stream-lined and user-friendly workflow to be adopted more broadly. Machine-mediated annotation tools, like CharaParser (Cui, 2012), are less time consuming, but, due to the high number of homonyms in biodiversity descriptions, these methods have a higher error rate.

## Challenges and future directions

Insect biodiversity research already utilizes ontologies (*e.g*., our hierarchical classification of Life), yet there remains some nescience or even skepticism regarding their application. Below we outline some misconceptions and real barriers to effective implementation of semantic approaches to phenotype representation. We also offer ideas for how to surmount these barriers, where possible, and describe a vision for the future of phenotype data in biodiversity research.

**Social challenges to “standardization”** A common misconception is that the ontology implementation is an attempt to force a research community to use a unified lexicon. Biodiversity research is extraordinarily heterogeneous, and each domain has its own history of language and “camps” of experts. Sociologically it’s impossible to get widespread agreement on terminology. An ontology *can* act as a controlled vocabulary, and best practices call for a single “preferred” term for each concept. The *sensu* model and glossary development described above, however, allow for the retention of personalized lexicons, while also facilitating communication through a standard set of concepts.

**Ontology development barriers** Several taxon-focused phenotype ontology development projects have been funded by the U. S. National Science Foundation (see awards DBI-0850223, DBI-0641025, DEB-0640053, for example), each employing a large staff of informaticians, postdocs, and other high-level personnel. Given this history, there is a perception that developing a new ontology, for hemipteran anatomy, for example, is an substantial challenge. Much of the heavy lifting—development of concepts, standards, tools, and documentation—has been done, however, which lowers barriers to future implementation. The Hymenoptera Anatomy Ontology, for example, could be cloned and modified with relative ease and applied to almost any hexapod taxon. This approach is being done for Neuroptera and Coleoptera, with relatively few resources. Documentation and examples of implementation are forthcoming.

**Ontology implementation barriers** Researchers who generate phenotype data, especially taxonomists, already document their science extensively, perhaps more so than any other domain in the life sciences. Adding yet another task, the generation of structured phenotype data, might further prolong the descriptive process at a time when biodiversity is rapidly disappearing. It is true that the current workflow is a bit complicated, requiring multiple applications and the use unfamiliar standards and syntax (see Mikó *et al*., 2015). There have been several calls for investment in phenotype data collection and analysis (see Deans *et al*. (2015) and references therein), however, and tool development is ongoing. The Gene Ontology project, for example, has implemented a templates-based approach (Dietze *et al*., 2014) that facilitates ontology implementation without technical knowledge of syntax and the ontologies themselves. We anticipate a relatively seamless, user-friendly integration of these approaches into biodiversity research in the near future.

**Phenotype complexity** Phenotypes can indeed be elaborate, and many natural language descriptions cannot be understood without figure references or the specimens in hand. Characters with graded quantifiers—*e.g*. “Scutellum strongly curved in lateral view (Fig. 6A)” *vs*. “Scutellum weakly curved in lateral view (Fig. 6B)” Baur *et al*. (2014)—and terms referring the putative evolutionary history, rather than pointing to an observable anatomical structure—*e.g*. “parossiculi fused” Mikó *et al*. (2013)—rely on non-textual references, *i.e*., images. The real meaning of these descriptions remains hidden behind the text (scutellum is more curved in species A than in species B, and an area of the integument is divided into two sclerites by a median conjunctiva) and cannot be translated adequately into semantic statements. Taxonomic descriptions are flooded with similar culprits that must receive only course-grained annotations (“scutellum curved”) or be revised (“median conjunctiva present”).

**Communicating primarily with semantic phenotypes** The default approach to describing phenotypes is currently natural language, and most researchers assume that the subtleties and extent of many characteristics can only adequately be described in prose. The learning curve for telegraphic natural language, however, might be at least as steep as the learning curve for formal description. We understand the aversion of domain experts against structured representations, especially with its unfamiliar cadence and heavy usage of parentheses, but these representations provide a much more objective and standardized way to describe phenotypes. Our current approach is to complement, rather than replace, natural language descriptions with semantic phenotype data.

**No clearinghouse for phenotype data** At this time there is no phenotypic analog to GenBank^®^, a well-developed, high demand database with a suite of tools that facilitate genetic research (Benson *et al*., 2005). Phenotype data for most insects currently reside in individual publications, distributed across more than 1,000 scientific journals (Deans *et al*., 2012b) or as files in different data repositories (the Dryad Digital Repository, for example). Researchers with an interest in a particular taxon must seek out these data to perform any analyses. For broader computation, one must rely on other methods (text mining), and this remains an opportunity for innovation.

**Reasoning challenges** Balhoff *et al*. (2013) demonstrate that for small data sets one can make inferences computationally, using existing semantic reasoners and in a short period of time, and this approach has utility for error-checking and small-scale discovery. Larger data sets, however, require substantial computing resources. There is a substantial community effort to improve reasoning algorithms, through grand challenges and other mechanisms, and we anticipate that useful, large-scale computation is on the horizon.

None of these barriers is insurmountable, and given the increasing interest in semantic approaches to phenotype representation we predict that this aspect of the taxonomic enterprise—describing the diversity of life—will remain as vibrant and relevant as ever.

## Acknowledgments

This material is based upon work supported by the U. S. National Science Foundation, under Grant Numbers DBI-0850223, DEB-0956049, DBI-1356381, and DEB-1353252. Any opinions, findings, and conclusions or recommendations expressed in this material are those of the author(s) and do not necessarily reflect the views of the National Science Foundation. James P. Balhoff (RTI International) provided invaluable comments to an earlier version of this manuscript.

